# The EEG spectral fingerprints of meditation and mind wandering differ between experienced meditators and novices

**DOI:** 10.1101/2021.07.06.451305

**Authors:** Julio Rodriguez-Larios, Eduardo A. Bracho Montes de Oca, Kaat Alaerts

## Abstract

Previous literature suggests that meditation training is associated with changes in participants’ experience during meditation practice. In this study, we assess whether putative differences in the experience of meditation between meditators and non-meditators are reflected in EEG spectral modulations. For this purpose, we recorded electroencephalography (EEG) during rest and two breath focus meditations (with and without experience sampling) in a group of 29 adult participants with more than 3 years of meditation experience and a control group of 29 participants without any meditation experience. Experience sampling in one of the meditation conditions allowed us to disentangle periods of breath focus from mind wandering (i.e. moments of distraction driven by task-irrelevant thoughts) during meditation practice. Overall, meditators reported a greater level of focus and reduced mind wandering during meditation practice than controls. In line with these reports, EEG spectral modulations associated to meditation and mind wandering also differed significantly between meditators and controls. While meditators (but not controls) showed a significant decrease in individual alpha frequency and amplitude and a steeper 1/f slope during meditation relative to rest, controls (but not meditators) showed a relative increase in individual alpha amplitude during mind wandering relative to breath focus periods. Together, our results show that the experience of meditation changes with training and that this is reflected in oscillatory and non-oscillatory components of brain activity.

## 1. Introduction

Meditation is an umbrella term involving heterogeneous techniques originating from different traditions (Analayo, 2019; Hanh, 1990; Kabat-Zinn, 2006; Krishnamurti, 2002). Nonetheless, we can find some common denominators across different meditation practices. On the one hand, an important part of meditation practices involve sustained attention to an object of meditation that usually entails body sensations (Matko et al., 2021; Matko & Sedlmeier, 2019). On the other hand, different meditations traditions instruct their practitioners to observe thoughts as they arise during meditation practice without attaching to them (Anālayo, 2019; Delorme & Brandmeyer, 2019; Lutz et al., 2015). Based on these commonalities, it can be argued that different meditation techniques train the same set of cognitive skills. In support of this idea, previous literature suggests that different meditation practices improve sustained attention capacity and the ability to detect and disengage from mind wandering episodes (i.e. task-irrelevant thoughts) (Chiesa et al., 2011; Delorme & Brandmeyer, 2019; Mrazek et al., 2012, 2013; Tang et al., 2007).

Electroencephalography (EEG) has been widely used to study the neural correlates of meditative states. Although different EEG signatures have been explored (Aftanas & Golocheikine, 2002; J. Gao et al., 2016; Irrmischer et al., 2018), most of previous studies focused on the amplitude of neural oscillations. In this way, some reviews have concluded that meditation is most commonly associated to increased amplitude of alpha oscillations (∼8-12 Hz) relative to resting state in both meditators and non-meditators (Lee et al., 2018; Lomas et al., 2015). However, these results have to be taken with caution since several studies have shown opposite effects (Aftanas & Golocheikine, 2001; Amihai & Kozhevnikov, 2014; Lehmann et al., 2012) or no changes at all in the alpha range (Braboszcz et al., 2017; Cahn et al., 2010).

It is has been suggested that (at least) part of the differences in EEG oscillatory activity between meditation and resting state could be related to mind wandering (Hinterberger et al., 2014; Lehmann et al., 2012; Rodriguez-Larios, Wong, et al., 2020). This hypothesis is motivated by self-reports from experienced meditators suggesting decreased mind wandering frequency and engagement during meditation practice relative to rest (Brandmeyer et al., 2019; Matko et al., 2021; Petitmengin et al., 2017). Consequently, several studies have assessed the EEG correlates of mind wandering during meditation practice (Braboszcz & Delorme, 2011; Brandmeyer & Delorme, 2018; Rodriguez-Larios & Alaerts, 2020; van Son et al., 2019). For this purpose, previous studies have used two types of experience sampling paradigms: i) self-caught experience sampling (i.e. participants press a button whenever mind wandering is noticed) and ii) probe-caught experience sampling (i.e. participants indicate whether they are mind wandering or not after they hear a bell sound) (Weinstein, 2018). The most consistent finding from studies using experience sampling and EEG during meditation practice is a relative decrease in alpha amplitude in mind wandering relative to states of focused attention (Braboszcz & Delorme, 2011; Brandmeyer & Delorme, 2018; Rodriguez-Larios & Alaerts, 2020; van Son et al., 2019). However, these findings directly contradict previous literature showing that mind wandering during different cognitive tasks is associated to a relative increase in alpha amplitude (Boudewyn & Carter, 2018; Compton et al., 2019; Gouraud et al., 2021; Groot et al., 2021; Jin et al., 2019, 2020).

Lapses of attention during different tasks are not only caused by mind wandering. In fact, moments of distraction can be also by caused by drowsiness, fatigue or ‘mind blanking’ (Andrillon et al., 2019, 2021). However, these other factors are typically not assessed directly in studies investigating the EEG correlates of meditation practice. This might be specially relevant in the case of drowsiness, as it has been shown to be highly correlated to mind wandering occurrence during meditation (Brandmeyer & Delorme, 2018). Since changes in drowsiness are also reflected by changes in EEG oscillatory activity (Jagannathan et al., 2018), it is possible that at least some of the identified oscillatory signatures of mind wandering during meditation practice are actually reflecting drowsiness. In support of this idea, a recent study has shown that mind wandering-related EEG oscillatory modulations were more pronounced for subjects who reported higher levels of drowsiness (Rodriguez-Larios & Alaerts, 2020).

The analytical approach adopted in previous studies could have also contributed to the lack of consensus regarding the EEG correlates of meditation practice and mind wandering. Thus, previous studies often estimated alpha power by defining its frequency band *a priori* (e.g. 8-12 Hz). This approach may however severely hinder consistency (e.g. in some studies alpha is divided in two sub-bands; Aftanas & Golocheikine, 2001, 2002; Saggar et al., 2012) and does not take into account inter-individual differences in alpha peak frequency (Klimesch, 1999b). More importantly, an *a priori* definition of frequency bands cannot disentangle whether changes in power are due to modulations in the peak amplitude of oscillatory activity, its peak frequency and/or the non-oscillatory components of the EEG spectrum (see Donoghue et al., 2021; Donoghue, Haller, et al., 2020; Kosciessa et al., 2020).

Neural oscillations are embedded within non-oscillatory / scale free neural activity that has a 1/f-like distribution (decreased power in higher frequencies)(Donoghue, Haller, et al., 2020). Recent evidence has shown that the commonly termed ‘1/f trend’ varies within subjects and is functionally relevant (Donoghue, Haller, et al., 2020; Kosciessa et al., 2021; Lendner et al., 2020; Voytek et al., 2015; Waschke et al., 2021). For example, the slope of the 1/f trend is thought to be reflective of excitation-inhibition balance in the brain (steeper slope is associated to greater inhibition) (R. Gao et al., 2017), thereby having important implications for cognition and consciousness (Kosciessa et al., 2021; Lendner et al., 2020; Ouyang et al., 2020). Crucially, it has been recently shown that changes in the 1/f trend of the EEG spectrum can conflate the estimation oscillatory activity (Donoghue, Dominguez, et al., 2020; Donoghue, Haller, et al., 2020). Since previous studies assessing the EEG correlates of meditation practice and mind wandering did not estimate nor control for the 1/f trend, it is possible that some of the inconsistencies in the literature are related to modulations in non-oscillatory activity.

In this study, we assessed whether the EEG correlates of meditation practice and mind wandering differ between experienced meditators and novices. For this purpose, we recorded EEG during rest and two breath focus meditations (with and without experience sampling) in a group of experienced meditation practitioners (N = 29) and a control group without any previous meditation experience (N = 29). Probe-caught experience sampling was used to disentangle periods of mind wandering from periods of breath focus during meditation practice. Unlike previous studies, we compared both oscillatory and non-oscillatory properties of the EEG signal between conditions (i.e. meditation vs rest; mind wandering vs breath focus) and groups (i.e. meditators vs controls). Specifically, four parameters of interest were assessed: i) power (2-30Hz) without an *a priori* definition of frequency bands, ii) the slope of the 1/f trend, iii) individual alpha power after controlling for the 1/f trend and iv) individual alpha frequency. In addition, we assessed the relationship between each of these parameters and drowsiness during meditation practice.

## 2. Methods

### 2.1 Participants

A total of 63 participants were recruited through social media and by directly contacting different meditation centers throughout Belgium. 5 participants had to be excluded due to technical problems during data acquisition. From the remaining 58 participants, 29 participants (12 males) had at least three years of experience with meditation practices (mean = 9.79 years; SD = 7.11) (i.e. meditators group). Meditators often reported to have experience with more than one meditation tradition. The three most reported traditions were Mindfulness (20 subjects), Zen (8 subjects) and Vipassana (9 subjects). The other 29 participants (14 males) had no previous experience with meditation practices (i.e. control group). The group of meditators and controls did not differ significantly in age (meditators, mean = 47.31, SD = 11.21; controls, mean = 47.13, SD = 13.93) (t-value (28) = 0.05; p-value = 0.95). Informed consent forms and study design were approved by the Social and Societal Ethics Committee (SMEC) of KU Leuven, in accordance with the Declaration of Helsinki (dossier no. G-2019 09 1747). Participants were compensated for their participation with 8 € per hour in addition to travel costs.

### 2.2 Design and task

EEG recordings were performed during three different conditions: rest (5 minutes), meditation (10 minutes) and meditation with probe-caught experience sampling (∼ 60 minutes) (for overview see **Figure 1A**). The experimental instructions were given in Dutch or English. During rest, participants were instructed to close their eyes, try to not to fall asleep, move as little as possible and let their minds wander in thought. For the meditation condition, the following instructions were used: *‘Sit in a comfortable posture that embodies dignity, keeping the spine straight and letting your shoulders drop. Close your eyes and allow your attention to gently align to the sensation of breathing. You can focus on the part of your body where you feel your breath most clearly (for example: nostrils, belly, chest…). Every time you notice that your mind has wandered off your breath, notice what it was that carried you away, and then gently bring your attention back to the sensations associated with your breath’* (Kabat-Zinn, 1990). In the meditation with experience sampling condition, participants were asked to follow the instructions appearing on a computer screen. First, they were asked to close their eyes and focus on their breath. After a period of 30 to 90 seconds, participants were presented with a bell sound and were required to open their eyes and report with a keyboard whether they were 1) focusing on their breath, 2) distracted by thoughts or 3) distracted by something else (sound, discomfort or other) (see **Figure 1D** for depiction). Regardless of the answer to the first question, participants had to report in the two following questions (scored on 7-point scales) their level of confidence on the previous question (ranging from ‘not confident at all’ (1) to ‘completely confident’ (7)) and their level of drowsiness (from ‘completely awake’ (1) to ‘falling asleep’ (7)). Then, some extra follow-up questions (also on 7-point scales) were asked depending on the answer to the first question. If participants answered ‘focusing on the breath’ in the first question, they were asked an extra question about the level of attention to the breathing (from ‘only superficially’ (1) to ‘full attention’ (7)). If participants answered ‘distracted by thoughts’ in the first question, they were asked three extra questions about the level of engagement in the thought (from ‘mostly observing’ (1) to ‘fully engaged’ (7)) and the level of visual and auditory components in the thought (from ‘not visual/auditory at all’ (1) to ‘completely visual/auditory’ (7)). Lastly, if participants answered ‘distracted by something else’ in the first question, they were asked in an extra question about what distracted them (three options: external sound, feeling of pain discomfort or something else). Participants performed a total of 40 trials in approximately 60 minutes.

**Figure 1.**
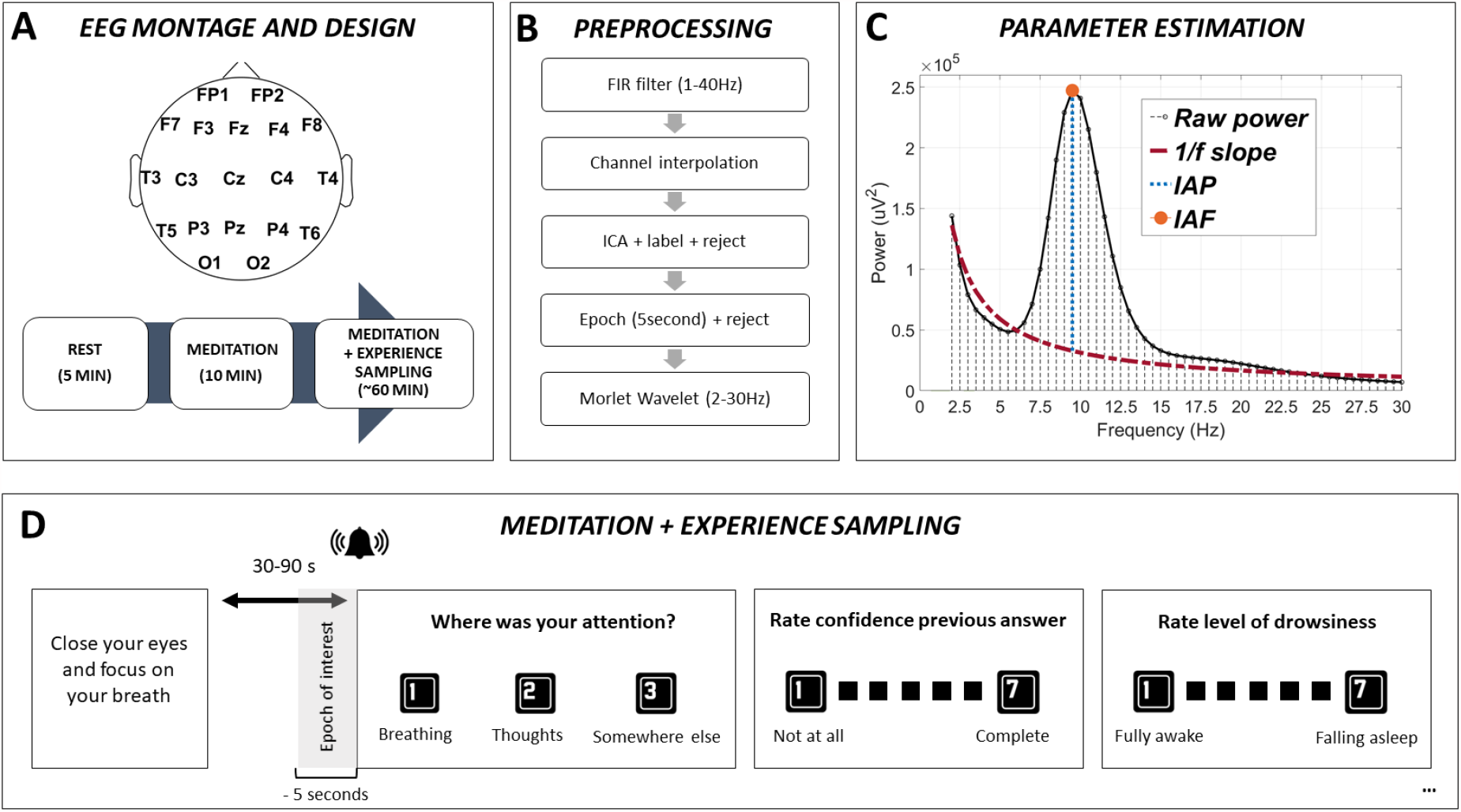
Design, EEG parameter estimation and task. A) Depiction of EEG montage and design. 19-channels EEG was acquired during three experimental conditions (rest, meditation and meditation with experience sampling) in a group of experienced meditators and a control group. B) Depiction of the pre-processing steps including: Finite impulse response (FIR) filter, channel interpolation, Independent Component Analysis (ICA), epoching and Morlet wavelet transform. C) Estimation of raw power, spectral 1/f trend slope, individual alpha power (IAP) and individual alpha frequency (IAF). D) Illustration of the meditation condition with experience sampling task. Participants were asked to close their eyes and focus on their breath. At random intervals between 30 and 90 seconds participants heard a bell sound and they were asked to report the location of their attentional focus (breathing, thoughts or something else). Then, several follow-up questions were asked for a better characterization of their experience (confidence, drowsiness…etc.). Participants performed 40 trials in ∼60 minutes. Only the 5 seconds before the bell sound (epoch of interest; see grey area) were used for further EEG analysis.

### 2.3 EEG data acquisition

Electroencephalography (EEG) recordings were performed using a Nexus-32 system and BioTrace software (V2018A1) (Mind Media, The Netherlands). Continuous EEG was recorded with a 19-electrode cap (plus two reference electrodes and one ground electrode) positioned according to the 10–20 system (see **Figure 1A**). Vertical (VEOG) and horizontal (HEOG) eye movements were recorded by placing pre-gelled foam electrodes (Kendall, Germany) above and below the left eye (VEOG) and next to the left and right eye (HEOG) (sampling rate of 2048 Hz). Skin abrasion and electrode paste (Nuprep) were used to reduce the electrode impedances during the recordings. The EEG signal was amplified using a unipolar amplifier with a sampling rate of 512 Hz. In the meditation with experience sampling condition, EEG recordings were synchronized to E-prime 2.0 using the Nexus trigger interface (Mind Media).

### 2.4 EEG analysis

Pre-processing was performed in MATLAB R2020b using custom scripts and EEGLAB (version 2019) functions (Delorme & Makeig, 2004) (see **Figure 1B** for overview). EEG data were first filtered between 1 and 40 Hz (function *pop_eegfiltnew*). Then noisy electrodes were detected automatically (function *clean_channels*) and interpolated (spherical interpolation implemented in function *pop_interp*). A mean of 1.40 channels (SD = 1.20) were interpolated. Then EEG data were re-referenced to the common average and independent component analysis (ICA) (function *pop_runica*) was performed. An automatic component rejection algorithm (i.e. *IClabel*) was employed to discard components associated to muscle activity, eye movements, heart activity or channel noise (Pion-Tonachini et al., 2019). In addition, components with an absolute correlation with EOG channels higher than 0.6 were also discarded. The mean number of rejected components was 1.37 (SD = 1.01). Data was then epoched in 5 second epochs (note that in the meditation condition with experience sampling these 5 seconds corresponded to the 5 seconds before the bell sound; see **Figure 1D**). Epochs with an absolute amplitude higher than 100 μV were excluded (mean percentage of epochs rejected = 4.5 %; SD = 6.07). For subsequent analyses, only the last 5 minutes of the uninterrupted meditation condition were considered (originally 10 minutes long) to make the number of epochs equal to the rest condition (5 minutes long).

The frequency spectrum between 2 and 30 Hz (0.5 Hz resolution) of each epoch was obtained using Morlet wavelet transform with a wave number of 6 cycles (as implemented in the function *BOSC_tf*; see Whitten et al., 2011). Four parameters of interest were extracted from the frequency spectrum of each electrode: power at each frequency (without removal of 1/f trend), 1/f trend slope, individual alpha power (IAP) (after removal of the 1/f trend) and individual alpha frequency (IAF) (see **Figure 1C** for depiction). Power at each frequency was extracted by squaring the real component (amplitude) of the convolution between the EEG signal and the family of wavelets. The 1/f slope of the spectrum was estimated by fitting a straight line in log–log space to the EEG frequency spectrum (excluding the 7 – 14 Hz range containing the alpha peak) using the *robustfit* function in MATLAB (for similar approaches see Caplan et al., 2015; Kosciessa et al., 2020; Watrous et al., 2018; Whitten et al., 2011). In order to estimate individual alpha, a find local maxima algorithm (function *findpeaks* in MATLAB R2020b) was employed to find a peak between 7 and 14 Hz above the estimation of the 1/f trend. Then, the amplitude of this peak relative to the 1/f trend (individual alpha power; IAP) and its frequency (individual alpha frequency; IAF) was calculated. The parameter estimation was performed in the average spectrum (across epochs) when comparing different conditions (i.e. meditation vs rest; mind wandering vs breath focus) and in an epoch by epoch basis to assess correlations within subjects (i.e. relation between drowsiness and each of the estimated parameters across different trials). Note that the average spectrum in the meditation with experience sampling condition (last 5 seconds before the bell sound) was estimated by performing a weighted average of the spectrum of different epochs, so epochs with a greater confidence rate had a greater weight in the average spectrum.

### 2.5 Statistical analysis

A cluster-based permutation statistical method (Maris & Oostenveld, 2007) was adopted to assess the significance of condition-related differences and trial by trial correlations in each EEG dependent variable. This statistical method controls for the type I error rate arising from multiple comparisons using a non-parametric Montecarlo randomization. First, cluster-level test statistics are estimated in the original data (comparing subjects between conditions or correlations against zero) and in several shuffled versions of the data (i.e. 10,000 random partitions). Cluster-level test-statistics are defined as the sum of t-values with the same sign across adjacent electrodes and/or frequencies that are above a specified threshold (i.e. 97.5th quantile of a T-distribution). Then, the cluster-level statistics from the original data and the null distribution emerging from the random partitions are compared. Cluster-corrected p-values are defined as the proportion of random partitions whose cluster-level test statistic exceeded the one obtained in the original (non-shuffled) data. Significance level for the cluster permutation test was set to 0.025 (corresponding to a false alarm rate of 0.05 in a two-sided test) (Maris & Oostenveld, 2007). Paired-samples t-test was chosen as the test statistic to compare conditions (meditation vs rest and mind wandering vs breath focus) and independent samples t-test was used to compare between groups (meditators vs controls). For self-reports and behavioural data, Wilcoxon rank sum test was employed to compare scores between groups and Wilcoxon signed rank test was used to compare scores within subjects (both implemented in MATLAB R2020b).

## 3. Results

### 3.1 Self-reports during meditation with experience sampling

On average, participants reported to be focusing on their breath in 63.69 % of the trials (SD = 16.07), distracted by thoughts in 23.57 % of the trials (SD = 11.88) and distracted by something else in 14.62 % of the trials (SD = 8.97). For the statistical analysis we focused on ‘breath focus’ and ‘distracted by thoughts’ (i.e. mind wandering) trials.

Relative to the control group, meditators reported a significantly higher proportion of ‘breath focus’ trials (z-value = 4.46; p <0.001) (see top panel **Figure 2A**) and a significantly lower proportion of ‘distracted by thoughts’ trials (z-value = -3.01; p = 0.0026) (see top panel of **Figure 2B**). In addition, the meditators group (relative to controls) reported a significantly higher mean level of attention in breath focus trials (z-value = 3.40, p < 0.001) (**Figure 2A** bottom panel) and a lower mean level of engagement in mind wandering trials (z-value = - 1.87, p = 0.06) (statistical trend) (**Figure 2B** bottom panel). Generally, reports of drowsiness were low in both meditators (mean = 1.92; SD = 1.02) and controls (mean = 2.14; SD = 1.08) and no significant differences between groups were evident (z-value = -1.4315, p = 0.15). Also, no significant group differences were revealed for ratings of confidence (z-value = 0.82, p = 0.40), visualization (z-value = 0.08, p = 0.93) or auditory components of mind wandering episodes (z-value = 1.35, p = 0.17).

**Figure 2.**
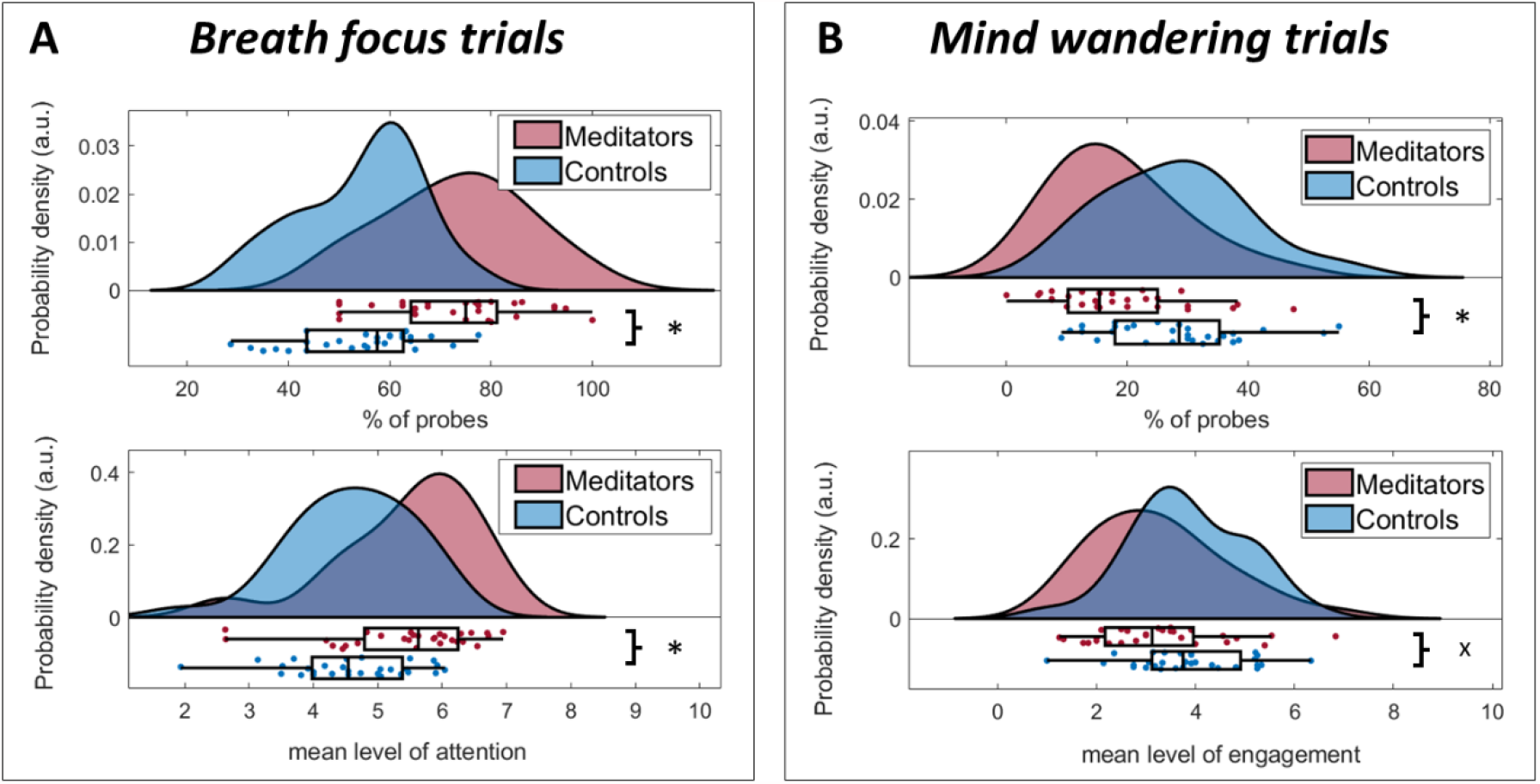
Self-reports during meditation with experience sampling. Individual data points and probability density distributions depicting the percentage of breath focus trials (A, top panel), the percentage of mind wandering trials (B, top panel), the mean level of attention in breath focus trials (A, bottom panel) and the mean level of engagement in mind wandering trials (B, bottom panel) for meditators (red) and control (blue) groups. Asterisks indicate significance (i.e. p<0.05) and ‘x’ indicates statistical tendency (p<0.1).

### 3.2 EEG spectral modulations associated to meditation relative to rest

We used cluster permutations statistics (see statistical analysis) to compare each of our EEG dependent variables (power at each frequency, 1/f slope, IAP and IAF; see **Figure 1C**) between conditions (meditation vs rest; paired samples t-test) and between groups (meditators vs controls; independent samples t-test).

#### Raw power

Experienced meditators showed a significant power decrease in the alpha/beta range (9 - 30Hz; widespread across electrodes) during meditation relative to rest (t_cluster_ = - 881.37; p_cluster_ < 0.001) (see **Figure 3D**). In contrast, the control group showed a trend-level high-beta (21 - 30 Hz) power increase in fronto-central electrodes during meditation (t_cluster_ = 91.86; p_cluster_ = 0.084) (see **Figure 3E**). A comparison of meditation-related changes between groups revealed that meditators showed a significantly more pronounced power decrease in the alpha/beta range (9-30Hz) (t_cluster_ = -666.56; p_cluster_ = 0.004) than controls (see **Figure 3F**).

**Figure 3.**
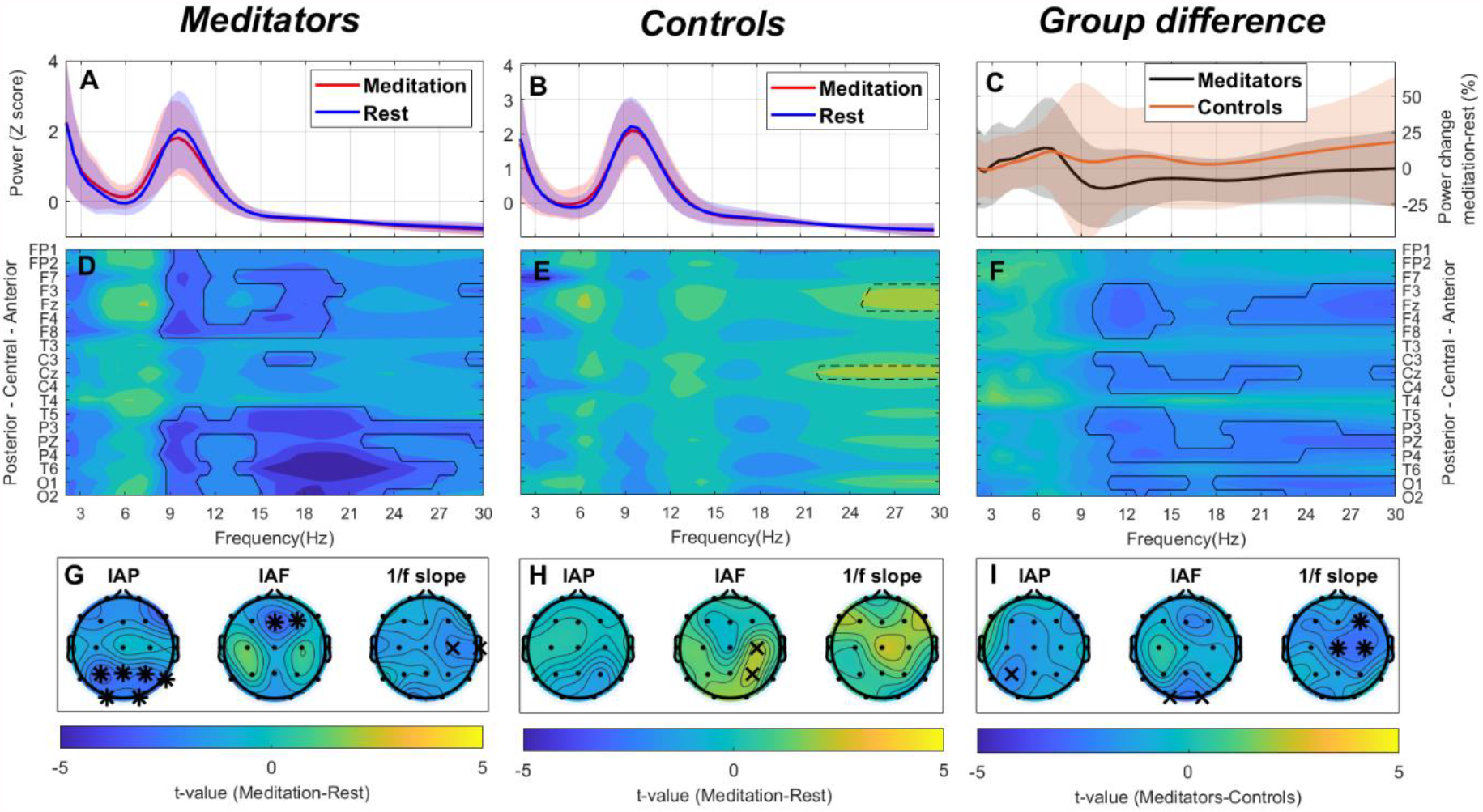
EEG spectral modulations associated to meditation in meditators and control groups. A-B) Average power spectrum (solid line) and standard deviation (shaded) in the 2-30 Hz range (across electrodes) during meditation (red) and rest (blue) in meditators (A) and the control (B) group. The power spectrum was z-scored (relative power) for visualization purposes. C) Mean power change (solid line) and standard deviation (shaded) in the 2-30 Hz frequency range (% change from rest; across electrodes) in meditators (black) and controls (orange). D-E) Matrix depicting the t-values resulting from comparing power at each frequency (x-axis; 2-30 Hz) and electrode (y-axis) between conditions (meditation vs rest) in the meditators (D) and control (E) group. The colours code for the direction of the differences between conditions (yellow = higher in meditation; blue = higher in rest). F) Matrix of t-values resulting from comparing power at each frequency (x-axis; 2-30 Hz) and electrode (y-axis) between groups during meditation. The colours code for the direction of the differences between groups (yellow = higher in meditators; blue = higher in controls). In all matrices, solid contours indicate significance at p<0.025 while dotted contours indicate statistical tendency (p<0.1). G-I) Topographical plots depicting differences in individual alpha power (IAP), individual alpha frequency (IAF) and the slope of the 1/f trend between conditions in each group (panel G for meditators and panel H for controls) and between groups (panel I). The colours code for the direction of the differences is the same as in the t-value matrices. Asterisks indicate statistical significance at p<0.025 whilst the symbol ‘x’ indicates statistical tendency (p<0.1).

#### Individual alpha power and frequency

Experienced meditators showed significant decrease in IAF in frontal electrodes (t_cluster_ = -5.42; p_cluster_ = 0.021) and significant decrease in IAP in parietal electrodes (t_cluster_ = -14.89; p_cluster_ = 0.010) during meditation relative to rest (see left and middle panels in **Figure 3G**). Instead, the control group only showed a trend-level IAF increase in central electrodes (t_cluster_ = 4.42; p_cluster_ = 0.039) during meditation (see left and middle panels in **Figure 3H**). A comparison of meditation-related changes between groups revealed that meditators showed a greater decrease in IAP in a parietal electrode (statistical trend) (t_cluster_ = -2.29; p_cluster_ = 0.072) and IAF in occipital electrodes (statistical trend) (t_cluster1_ = -2.49; p_cluster1_ = 0.093; t_cluster2_ = -2.47; p_cluster2_ = 0.096) than controls (see left and middle panels in **Figure 3I**).

#### 1/f slope

Experienced meditators showed a steeper 1/f slope (more negative values) during meditation relative to rest (t_cluster_ = -4.23; p_cluster_ = 0.057) (statistical trend) (see right panel in **Figure 3G**). Instead, controls showed a non-significant flattening (more positive values) of the slope during meditation (t_cluster_ = 2.54; p_cluster_ = 0.10) (see right panel in **Figure 3H**). Groups differed significantly in meditation-related changes in the 1/f slope (steeper during meditation for meditators only) (t_cluster_ = -7.76; p_cluster_ = 0.016) (see right panel in **Figure 3I**).

In summary, experienced meditators and controls differed significantly in meditation-related EEG spectral changes, and these changes encompassed both oscillatory and non-oscillatory components. Specifically, only experienced meditators showed reduced power in the alpha/beta band, reduced individual alpha power/frequency, and a steeper 1/f slope during meditation relative to rest.

### 3.3 EEG spectral modulations associated to mind wandering during meditation practice

We used the same cluster permutation procedure to compare each of our EEG dependent variables (power at each frequency, 1/f slope, IAP and IAF; see **Figure 1C**) between mind wandering and breath focus periods identified during the meditation with experience sampling condition (see **Figure 1D**). In addition, we also compared mind wandering-related changes in each of our dependent variables between meditators and the control group.

#### Raw power

Experienced meditators showed a trend-level decrease in theta band power (3 - 6 Hz) in frontal electrodes during mind wandering relative to breath focus (t_cluster_ = 67.72; p_cluster_ = 0.078) (see **Figure 4D**). In contrast, controls showed a significant increase in alpha/beta (10.5 - 25 Hz) power during mind wandering across electrodes (t_cluster_ = 485.32; p_cluster_ = 0.0064) (see **Figure 4E**). Significant differences between groups in mind wandering-related changes were found in both theta and alpha/beta bands (3-25 Hz range) (t_cluster_ = - 1192.99; p_cluster_ = 0.0017) (see **Figure 4F**).

**Figure 4.**
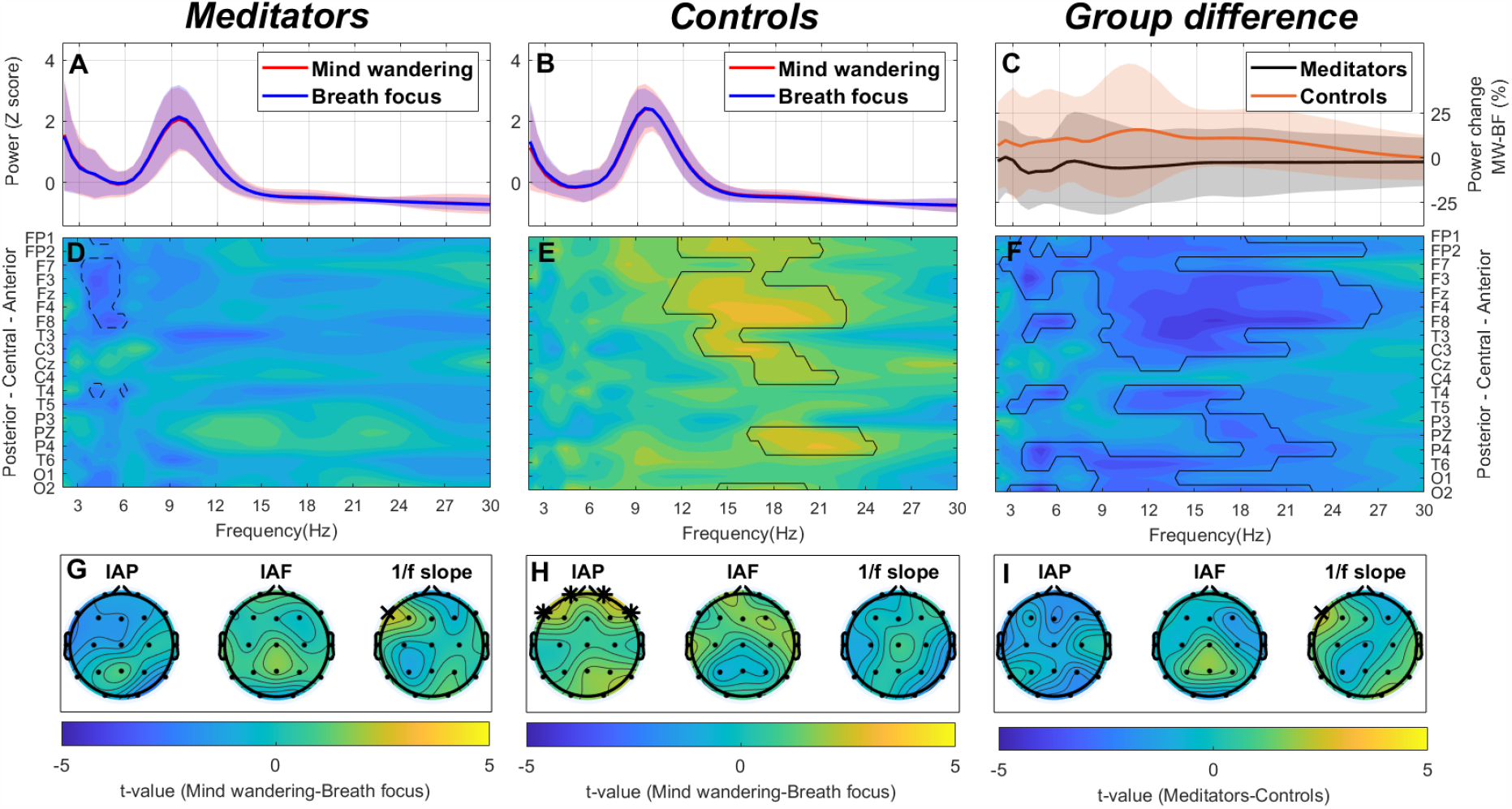
EEG spectral modulations associated to periods of mind wandering relative to periods breath focus in meditators and control groups. A-B) Average power spectrum (solid line) and standard deviation (shaded) in the 2-30 Hz range across electrodes during mind wandering (red) and breath focus (blue) in meditators (A) and control (B) groups. The power spectrum was z-scored (relative power) for visualization purposes. C) Mean power percentage change (mind wandering – breath focus) (solid line) and standard deviation (shaded) in the 2-30 Hz frequency range across electrodes in meditators (black) and controls (orange). D-E) Matrix depicting the t-values resulting from comparing power at each frequency (x-axis; 2-30 Hz) and electrode (y-axis) between conditions (mind wandering vs breath focus) in the meditators (D) and control (E) group. The colours code for the direction of the differences between conditions (yellow = higher during mind wandering; blue = higher during breath focus). F) Matrix of t-values resulting from comparing power at each frequency (x-axis; 2-30 Hz) and electrode (y-axis) between groups during mind wandering (relative to breath focus). The colours code for the direction of the differences between groups (yellow = higher in meditators; blue = higher in controls). In all matrices, solid contours indicate significance at p<0.025 while dotted contours indicate statistical tendency (p<0.1). G-I) Topographical plots depicting differences in individual alpha power (IAP), individual alpha frequency (IAF) and the slope of the 1/f trend between conditions in each group (panel G for meditators and panel H for controls) and between groups (panel I). The colours code for the direction of the differences in each panel is the same as in the t-value matrices. Asterisks indicate statistical significance at p<0.025 whilst the symbol ‘x’ indicates statistical tendency (p<0.1).

#### Individual alpha power and frequency

No differences in experienced meditators were found. Only the control group showed a significant increase of IAP in frontal electrodes during mind wandering relative to breath focus (t_cluster_ = 9.03; p_cluster_ = 0.019) (see left panel in **Figure 4H**). However, group differences in mind wandering-related IAP modulations did not reach statistical significance.

#### 1/f slope

Experienced meditators showed a trend-level flattening (more positive value) of the 1/f slope in a left frontal electrode in mind wandering relative to breath focus conditions (t_cluster_ = 2.62; p_cluster_ = 0.097) (see right panel in **Figure 4G**) while no differences were found in controls. This trend was also reflected in the group comparison (t_cluster_ = 2.56; p_cluster_ = 0.092) (see right panel in **Figure 4I**).

In summary, experienced meditators and controls differed significantly in mind wandering-related changes in the EEG spectrum during meditation practice. While meditators showed a trend-level decrease in theta band power and trend-level flattening of the 1/f slope during mind wandering relative to breath focus, controls showed a significant increase in alpha/beta band power and IAP.

### 3.4 EEG spectral modulations associated to drowsiness during meditation

In addition to the assessment of condition-related differences, we assessed the relationship between trial-by-trial variations in each of our dependent variables and trial-by-trial variations in self-reported drowsiness during meditation practice. For this purpose, Spearman rank order correlations were estimated for each of the dependent variables (power at each frequency, 1/f slope, IAP and IAF; see **Figure 1C**) per electrode and subject. Then, we assessed the significance of these correlations at group level using the described cluster permutation test (Maris & Oostenveld, 2007) against zero (one sample t-test). In addition, we assessed potential group differences in the obtained correlations using the same cluster permutation test but with independent samples t-test as the test statistic.

#### Raw power

Drowsiness was positively correlated to theta/low-alpha band power (3 – 8 Hz) in both experienced meditators (t_cluster_ = 138.10; p_cluster_ = 0.059) (statistical trend; see **Figure 5D**) and controls (t_cluster_ = 399.10; p_cluster_ = 0.016) (see **Figure 5E)**. No significant differences were found between groups.

**Figure 5.**
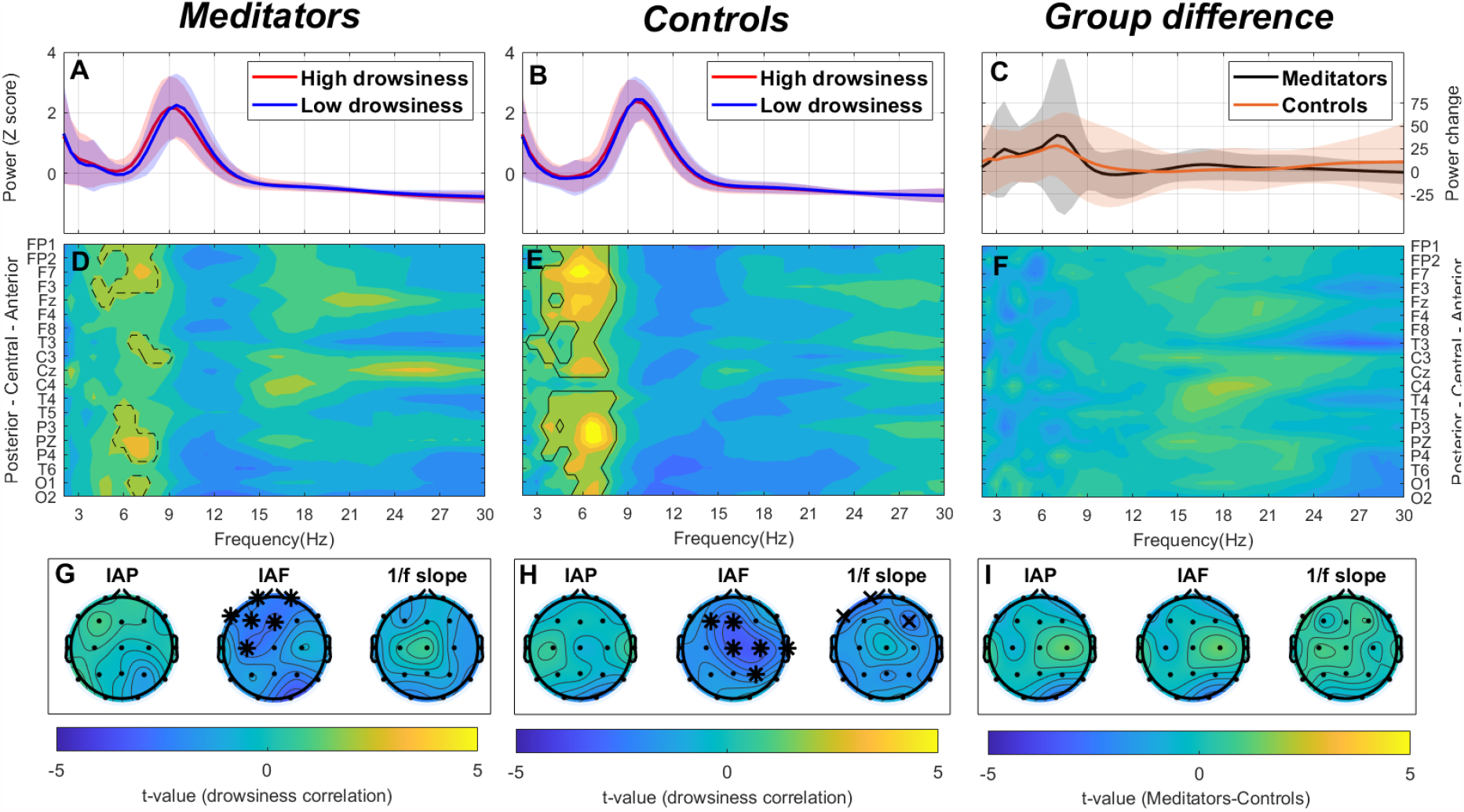
EEG spectral modulations associated to drowsiness during meditation with experience sampling across groups. A-B) Average power spectrum (solid line) and standard deviation (shaded) in the 2-30 Hz range across electrodes during high drowsiness (red) and low drowsiness (blue) in meditators (A) and control (B) groups. The power spectrum was z-scored (relative power) and a median split was performed based on drowsiness for visualization purposes. C) Mean power percentage change (high – low drowsiness) (solid line) and standard deviation (shaded) in the 2-30 Hz frequency range across electrodes in meditators (black) and controls (orange). D-E) Matrix depicting the t-values resulting from the statistical assessment of the correlation between power at each frequency (x-axis; 2-30 Hz) and electrode (y-axis) and drowsiness in the meditators (D) and control (E) group. The colours code for the direction of the correlation with drowsiness (yellow = positive correlation; blue = negative correlation). F) Matrix of t-values resulting from comparing the correlation with drowsiness at each frequency (x-axis; 2-30 Hz) and electrode (y-axis) between groups. The colours code for the direction of the differences between groups (yellow = higher in meditators; blue = higher in controls). In all matrices, solid contours indicate significance at p<0.025 while dotted contours indicate statistical tendency (p<0.1). G-I) Topographical plots depicting the correlations between individual alpha power (IAP), individual alpha frequency (IAF) and the slope of the 1/f trend and drowsiness in each group (panel G for meditators and panel H for controls) and the differences between groups (panel I). The colours code for the direction of the correlations and group differences is the same as in the t-value matrices. Asterisks indicate statistical significance at p<0.025 whilst the symbol ‘x’ indicates statistical tendency (p<0.1).

#### Individual alpha power and frequency

Drowsiness was negatively correlated to IAF in both experienced meditators (t_cluster_ = -14.74; p_cluster_ = 0.004) (see middle panel in **Figure 5G**) and controls (t_cluster_ = -15.98; p_cluster_ = 0.006) (see middle panel in **Figure 5H**). No significant differences were found between groups.

#### 1/f slope

A steeper 1/f slope was associated to increased drowsiness in controls (statistical trend) (t_cluster_ = -5.19; p_cluster_ = 0.04) (see **Figure 5H** right panel). Although meditators showed correlations in the same direction, it did not reach statistical significance (t_cluster_ = -2.17; p_cluster_ = 0.21) (see **Figure 5G** right panel). No significant differences were found between groups.

In summary drowsiness during meditation practice was associated to increased power in high-theta/low-alpha band, reduced individual alpha peak frequency and a steeper 1/f slope. Importantly, no significant differences were found between groups in drowsiness-related EEG spectral modulations.

## 4. Discussion

This study aimed to clarify previous inconsistencies in the literature about the EEG spectral modulations associated to meditation and mind wandering in participants with and without meditation experience. For this purpose, we compared meditation/mind wandering-related EEG spectral modulations between a group of experienced meditators and a control group without any meditation experience. Overall, meditators reported reduced mind wandering and greater level of focus during meditation practice than the control group. Crucially, group differences in the experience of meditation were accompanied by group differences in meditation/mind wandering-related EEG spectral modulations. On the one hand, only the group of experienced meditators showed significant EEG spectral changes between the meditation and rest. Specifically, meditators showed a power decrease in alpha/beta band, a decrease in individual alpha frequency and a steeper 1/f slope during meditation. On the other hand, only participants without meditation experience (i.e. control group) showed significant EEG spectral modulations associated to mind wandering during meditation practice. In particular, the control group showed a significant power increase in alpha/beta band during mind wandering relative to periods of breath focus. In addition, further analyses revealed that the EEG spectral modulations associated to drowsiness during meditation practice (i.e. reduced individual alpha frequency and greater power in theta/low-alpha band) did not differ significantly between meditators and the control group.

In this study, we show that, oscillatory activity in the alpha range decreases in power and frequency during breath focus meditation relative to rest in experienced meditators only. Our results are in line with other studies reporting a relative decrease in alpha power (Aftanas & Golocheikine, 2001; Amihai & Kozhevnikov, 2014; Lehmann et al., 2012) and frequency (Aftanas & Golocheikine, 2002; Irrmischer et al., 2018; Rodriguez-Larios, Faber, et al., 2020; Saggar et al., 2012; Takahashi et al., 2005; Yamamoto et al., 2006) during meditation. However, it is important to underline that previous literature reviews have concluded that meditation is most commonly associated to relative increases in alpha power when compared to rest and other control conditions (Lee et al., 2018; Lomas et al., 2015). It is possible that this inconsistency relates to the fact that previous reviews included meditation techniques that significantly differ from breath focus meditation (e.g. mantra recitation, visualizations…etc) and/or included subjects without meditation experience (Lee et al., 2018; Lomas et al., 2015). Another possibility is that the lack of consensus in the literature is related to the adopted analytical approach. In this regard, since previous studies estimated power in an *a priori* definition of the alpha band (i.e. ∼8 – 13 Hz) without peak detection nor control for the 1/f trend of the spectrum, we cannot rule out the possibility that their results are a mix of changes in oscillatory power, frequency and/or the 1/f slope (Donoghue, Dominguez, et al., 2020; Donoghue, Haller, et al., 2020).

Experience sampling during meditation practice revealed that only non-meditators showed a significant increase in alpha band power during mind wandering relative to breath focus (see **Figure 4E** and left panel of **Figure 4H**). Since less mind wandering engagement was reported in meditators relative to non-meditators, it can be speculated that for mind wandering to be associated to a relative increase in alpha power, a certain level of engagement in mind wandering (i.e. attachment to thoughts; Lutz et al., 2015) is necessary. In this way, it is important to underline that although mind wandering has been consistently associated to increased alpha power in a wide variety of cognitive tasks (Boudewyn & Carter, 2018; Compton et al., 2019; Gouraud et al., 2021; Groot et al., 2021; Jin et al., 2019, 2020), three previous studies using experience sampling during meditation practice with non-experienced practitioners showed the opposite pattern of results (i.e. reduced alpha power during mind wandering) (Braboszcz & Delorme, 2011; Rodriguez-Larios & Alaerts, 2020; van Son et al., 2019). On the one hand, two of these previous studies employed a self-caught experience sampling paradigm (i.e. participants are instructed to press a button whenever they realize that they are mind wandering) (Braboszcz & Delorme, 2011; van Son et al., 2019) instead of a probe-caught experience sampling paradigm (i.e. participants are instructed to report whether they are mind wandering after they hear a bell sound). Hence, it is possible that self-caught and probe-caught mind wandering are associated to different EEG spectral modulations. In fact, periods of mind wandering identified in self-caught paradigms also include other components such a meta-awareness and motor preparation that have been previously associated to alpha suppression (Deiber et al., 2012; van Driel et al., 2012). Nonetheless, it is important to note that our results are also inconsistent with our previous study, in which we did employ a probe-caught experience sampling paradigm and still reported reduced alpha power during mind wandering relative to breath focus (Rodriguez-Larios & Alaerts, 2020). In this regard, we speculate that the lapses of attention identified in our previous study were reflecting drowsiness and/or hypnagogic states instead of mind wandering *per se*. In support of this idea, we showed in Rodriguez-Larios & Alaerts (2020) that participants reported relatively high levels of drowsiness and that inter-individual differences in drowsiness levels were significantly correlated to both mind wandering occurrence and mind wandering-related EEG spectral modulations. In addition, the spectral profile identified for mind wandering in Rodriguez-Larios & Alaerts (2020) looks highly similar to the one that we associated to drowsiness in the current study (see **Figure 5E** this paper and Figure 2 in Rodriguez-Larios & Alaerts (2020)). Together, our results support the idea that mind wandering is associated to a relative increase in alpha power in different cognitive tasks (Boudewyn & Carter, 2018; Compton et al., 2019; Gouraud et al., 2021; Jin et al., 2019, 2020) and we hypothesize that previous inconsistencies in the literature could be explained by confounders such as motor preparation, meta-awareness and drowsiness.

The exact role of alpha oscillations in cognition and consciousness is still debated. The dominant model proposes that alpha oscillations exert top-down control by disinhibiting task-relevant neural populations and inhibiting task-irrelevant ones (Bonnefond & Jensen, 2012; Jensen et al., 2012; Jensen & Mazaheri, 2010). While alpha power has been consistently linked to (local) inhibition, there is no consensus about the meaning of changes in alpha peak frequency. In this regard, it is still unclear whether condition-related changes in alpha frequency reflect modulations in activity originating from the same or from different networks (Haegens et al., 2014; Mierau et al., 2017). In this way, if different alpha frequency ranges reflect the activity of different networks (Barzegaran et al., 2017; Benwell et al., 2019; Klimesch, 1999a; Sadaghiani et al., 2010) the here reported modulations in alpha frequency during meditation relative to rest could be reflective of a differential inhibition of different networks. In this regard, previous literature has shown that during concentrative meditation (relative to rest) the activity of the default mode network is supressed whilst regions involved in self-monitoring, sensory perception and cognitive control (i.e. task positive network) are more active (Brewer et al., 2011; Mahone et al., 2018; Pagnoni et al., 2008; Winter et al., 2020). Therefore, it can be speculated that lower alpha band (∼ 7 - 9 Hz) inhibits task-irrelevant regions during meditation (e.g. default mode network) while upper alpha (∼ 10 – 12 Hz) inhibits task-irrelevant regions during rest (e.g. task positive network). In the same line, changes in alpha power in the absence of changes in individual alpha frequency (mind wandering in non-experienced practitioners; see **Figure 4H**) could be interpreted as a change in activity or connectivity of a specific network (e.g. the default mode network) (Chapeton et al., 2019; Lobier et al., 2018; Samogin et al., 2019, 2020). In order to empirically assess these speculations, further research is needed to test whether i) different brain networks oscillate at different alpha frequencies and ii) whether the level of activity or connectivity within a network modulates alpha power at EEG sensor level. For this purpose, it is imperative to identify the spectral profile of different networks in the future by either combining EEG with intracranial electrodes (Fahimi Hnazaee et al., 2020) or through high density EEG and source localization (Samogin et al., 2019, 2020).

In addition to the reported changes in the alpha range, we also found significant group differences in meditation/mind wandering-related spectral modulations in the theta (∼ 3 -7 Hz) and beta (∼ 12 - 30 Hz) bands. Recent evidence has shown that changes in the 1/f slope can especially conflate estimations of oscillatory power in theta and beta ranges, where clear spectral peaks (which is the most robust signature of oscillatory activity) are not easily observed (Donoghue, Dominguez, et al., 2020). However, our results show that modulations in theta and beta band power are not always accompanied by changes in the slope of the 1/f trend. In particular, we found a significant beta power increase (∼ 12 - 24 Hz) during mind wandering relative to breath focus periods in the control group that was not accompanied by a significant change in the 1/f slope (see **Figure 4E** and right panel of **Figure 4H**). Therefore, it is possible that the here reported changes in beta band power do reflect oscillatory activity but that due to a low signal to noise ratio and/or the non-stationarity of the signal (Nikulin et al., 2011; van Ede et al., 2018; Vidaurre et al., 2018) this is not reflected as a clear peak in the frequency spectrum. If this is the case, there are at least two possible interpretations of a relative increase in beta oscillatory power during mind wandering in the context of meditation practice. On the one hand, in the light of recent accounts on the role of beta oscillations in cognition (Spitzer & Blankenburg, 2012; Spitzer & Haegens, 2017), it can be speculated that a relative increase in beta oscillatory activity during mind wandering is associated to the reactivation of content-specific cortical representations (i.e. those associated to memories, fantasies, plans…etc.). This interpretation would be in line with previous studies showing reduced beta oscillations in highly experienced meditators during meditative states in which the emergence of thoughts is expected to be minimized (Dor-Ziderman et al., 2013, 2016). Another possibility is that power changes in the beta band are a consequence of non-sinusoidal alpha oscillations originating from the sensorimotor cortex, which are known to also be reflected in the beta range (Angelini et al., 2018; Fox et al., 2016; Schaworonkow & Nikulin, 2019). In line with this idea, previous studies have shown that sensorimotor alpha oscillations (and its corresponding beta component) are suppressed during focused attention to body sensations in the context of meditation practice (Kerr et al., 2011, 2013). Given the putative inhibitory function of alpha oscillations (Bonnefond & Jensen, 2012; Jensen et al., 2012; Jensen & Mazaheri, 2010), it can be hypothesized that alpha/beta power increases during mind wandering relative to breath focus are due to a greater inhibition of the sensorimotor cortex when subjects are not fully paying attention to body sensations associated to the breath.

Unlike meditation and mind wandering, the EEG spectral modulations associated to drowsiness did not differ significantly between meditators and non-meditators. In line with previous studies (Broughton & Hasan, 1995; Cantero et al., 2002), drowsiness was associated to an increase in theta/low-alpha band power and a decrease in individual alpha frequency (IAF) (see **Figure 5**). In addition, drowsiness was also associated to a steeper 1/f slope (statistical tendency in control group; see **Figure 5H** right panel). Interestingly, some of the EEG spectral modulations associated to drowsiness (steeper 1/f slope and reduced IAF) were also identified when comparing meditation and rest conditions in experienced meditation practitioners (see **Figure 3G**). In this way, both a relative decrease in IAF and a steeper 1/f slope have been previously associated to reduced arousal levels (Cantero et al., 2002; Kosciessa et al., 2021; Lendner et al., 2020; Mierau et al., 2017). Since a relative decrease in arousal can be expected during both drowsiness and a state of relaxation during meditation practice (Jagannathan et al., 2018; Matko et al., 2021; Petitmengin et al., 2017), we speculate that steeper 1/f slope and reduced IAF during meditation in experienced practitioners are reflective of reduced arousal due to relaxation. In this way, it is important to note that the topography of drowsiness and meditation-related changes in IAF and the 1/f slope seem to differ (see **Figure 3G** and **Figure 5G-H**). Therefore, it is possible that drowsiness and relaxation are characterized by reduced IAF and a steeper 1/f slope in different regions. In this line, future experience sampling studies should be warranted to specifically address how the EEG spectral correlates of drowsiness and relaxation differ.

To our knowledge, there are two main limitations of the study that need to be addressed. The first limitation is related to the relatively low number of EEG electrodes employed in this study (i.e. 19), which only allowed us to draw conclusions about EEG spectral changes at a sensor level. In this way, high-density EEG and source localization analysis (Michel & Brunet, 2019; Samogin et al., 2019) should be warranted in the future to reveal the sources of the here reported EEG changes. Furthermore, source localization analysis might also allow to identify robust oscillatory activity in other bands in addition to alpha (i.e. spectral peaks in the theta and beta bands) (Dasari et al., 2017; Grandchamp et al., 2012). The second limitation is related to the use of experience sampling paradigms to study mind wandering. In this context, experience sampling paradigms entail a trade-off between the length of the inter-probe time interval (it should be long enough so mind wandering actually occurs) and sufficient number of probes (so sufficient mind wandering epochs can be identified). In this way, if the duration of the experiment is extended to increase both the time interval between probes and the number of probes, participants are more likely to report fatigue and drowsiness. Consequently, experience sampling paradigms tend to identify a relatively low number of mind wandering periods, above of all, when participants can also report lapses of attention unrelated to mind wandering. In our case, an average of 23 % of the probes were classified as mind wandering (across groups), which corresponds to an average of 9 mind wandering epochs per participant. In order to overcome this limitation, future studies could perform multiple (mobile) EEG recording sessions in the same subjects (Reiser et al., 2020) thereby allowing the identification of a greater number and variety of mind wandering periods.

In summary, this study compared self-reports during meditation and meditation-related EEG spectral modulations between a group of experienced meditators and a control group without previous meditation experience. Self-reports revealed that meditators showed a greater level of focus and reduced mind wandering frequency and engagement than controls during meditation practice. In line with these reports, the EEG spectral modulations associated to meditation (relative to rest) and mind wandering (during meditation) also differed significantly between meditators and controls. While meditators (but not controls) showed a decrease in alpha/beta band power, reduced individual alpha frequency and amplitude and a steeper 1/f slope during meditation relative to rest, controls (but not meditators) showed a relative increase in alpha/beta band power and individual alpha power during mind wandering relative to breath focus. Based on these results, we conclude that the experience of meditation changes with meditation training and that this is reflected in oscillatory and non-oscillatory components of brain activity.

## 5. Additional information

## Acknowledgements

This work was supported by the Branco Weiss fellowship of the Society in Science–ETH Zurich, Grants from the Flanders Fund for Scientific Research (FWO G079017N and G046321N) and the European Varela Awards (Mind & Life Europe). We also would like to thank all the volunteers that participated in the study.

## Competing interests

The authors declare no competing interests.

## Author contributions

J.R. and K.A. conceptualized the study and acquired the funding. E.B. organized and performed the data collection. J.R. performed the analysis and wrote the manuscript. J.R., K.A. and E.B. revised the manuscript and approved the final version.

## Data availability

Raw EEG data and MATLAB code will be publicly available in the Open Science Framework webpage (see https://osf.io/3uszv/?view_only=d41ddd2200e642cf9992a016cb739b90).

## Notes

### Competing Interest Statement

The authors have declared no competing interest.

